# Changes in free-roaming dog population demographics and health associated with a catch-neuter-vaccinate-release program in Jamshedpur, India

**DOI:** 10.1101/2025.01.05.631379

**Authors:** Lauren Smith, Tamara Kartal, Sanjay Rawat, Amit Chaudhari, Ashok Kumar, Rajesh Kumar Pandey, Rupert J. Quinnell, Lisa Collins

## Abstract

India’s large free-roaming dog populations contribute to significant human health, environmental, and social challenges. Population management strategies, such as capture-neuter-vaccinate-return (CNVR), aim to reduce dog numbers, improve their welfare, and reduce human-animal conflict. The Humane Society International, in partnership with the Animal Health Foundation, implemented a CNVR program in Jamshedpur, neutering and vaccinating over 20,000 dogs. This study evaluates the impact of this program on dog health, population structure and size. The study areas encompassed 10 sites within the Jamshedpur Metropolitan Region, including both intervention sites where CNVR was directly applied and sites without direct intervention. Data was collected from May 2014 until December 2018, including bi-annual street surveys, as well as clinical data from the dogs captured and treated. We fit logistic regression, negative binomial, and binomial mixed effects models to assess changes in dog population characteristics, health, and reproductive conditions over time in relation to the CNVR intervention. We found that, over time, the CNVR program significantly reduced the probability of dogs entering the clinic with mange, transmissible venereal tumours, and pregnant. Street surveys showed an increase in sterilised dogs, with higher proportions observed in CNVR-treated sites, although the counts of dogs observed increased overall. The age-structure of free-roaming dogs remained stable over time. In CNVR-treated areas, the probability of observing lactating female dogs decreased, whereas it increased in untreated sites. This work contributes to the growing body of knowledge investigating the impact of dog population management interventions. Continued monitoring and evaluation of CNVR programs are required to identify optimal coverage required to reduce population size effectively.

## 1. Introduction

The worldwide population of domestic dogs (*Canis familiaris*) is estimated to be between 700 million to 1 billion (1,2), a high proportion of which are “free-roaming” for all, or at least part of, the day (1). Free-roaming dogs can roam and reproduce without restriction, with possible consequences for public health (3–5), wildlife conservation (6–8), and livestock losses (9–13). Free-roaming dogs may also experience high mortality and poor welfare (14–17). India has high densities of free-roaming dogs, with estimates in rural areas as high as 719 dogs per km^2^ (18). Consequently, the country faces serious public health, environmental, and social challenges. There are an estimated 20 million dog bites in India per year (approximately 150 bites per 10,000 people) (19), and victims often require medical attention or post-exposure rabies prophylaxis. Dogs are responsible for 99% of human-rabies transmissions and, although the true rabies burden in India is unknown, it is estimated to result in 18,000 to 20,000 human deaths per year (approximately 35% of the worldwide rabies burden) (19,20). Free-roaming dogs are also responsible for the killing of livestock, such as sheep, goats, and donkeys, contributing to substantial economic losses, and exacerbating human-wildlife conflict, as predation by dogs can be mistaken for predation by wolves or snow leopards (9,21). There are also beneficial aspects of the free-roaming dog population: dogs may provide companionship, guard livestock, and reduce vermin.

In India, the free-roaming dog population is dependent on humans as a source of food, either directly by households providing food, or indirectly, for example, through improper refuse disposal (22). It is estimated that free-roaming dogs in India have a short lifespan of less than four years (23), and that only 18% of puppies survive to one year old (24,25). Birth rates are high and almost 50% of female dogs are estimated to become pregnant in any given year (23). These high mortality and birth rates lead to high population turnover, which makes maintaining high vaccination coverages challenging for infectious disease control, such as rabies (26).

Dog population management can reduce the population size or the risks associated with free-roaming dogs, and improve the health and welfare of the dog population. Dog population management methods can include culling, sheltering, fertility control, reducing the carrying capacity of the population (e.g. by reducing the availability of resources (22)), and encouraging responsible ownership practices (27). In India, national law requires that street or community owned (i.e. free-roaming) dogs are managed through the Animal Birth Control Rules of 2023 (28). This law stipulates that free-roaming dogs must be managed through fertility control, where dogs are caught, neutered, vaccinated against rabies (i.e. capture-neuter-vaccinate-return; CNVR), and returned to the same area where they were captured, once they complete the mandatory post-operative care period of three days.

Several studies have attempted to evaluate the impact of dog population management strategies, reviewed by (27). In India, the reported impact of fertility control on the health and welfare of free-roaming dogs varies, with both positive (29) and negative (29,30) impacts. This includes reports of improved body condition scores (31,32) and reduced presence of visible injuries (32), and both increased and decreased prevalence of pathogens and visible skin conditions (32). These effects have been attributed to a lack of sex hormones, which may reduce sexual competition and mate-seeking behaviour, which in turn can lead to decreased aggression (33) and improved body conditions (34,35). Fertility control programs, such as CNVR, regularly involve vaccination and antiparasitic treatments, which may also improve health and welfare. Fertility control programs aim to reduce the population size by reducing the population’s birth rate. Both no effect (36) and reductions in dog population size (37,38) have been reported following fertility control programs in India. In turn, a reduction in population size and increase in rabies vaccination following neutering and vaccination campaigns in free-roaming dog populations may reduce human bite and rabies cases. Reductions in both human bite cases (39) and human rabies cases (37) have been reported following neutering and vaccination campaigns in India. These impacts have been associated with varying intervention coverage (e.g. neutering coverage) and lengths of management.

Understanding the impact of CNVR is important for informing future dog population management efforts, so that we can optimise interventions and reduce the risks to humans and other animals that are associated with free-roaming dogs. The Humane Society International (HSI) is one of the largest animal protection and welfare organisations that works on almost all types of animal welfare issues. Through effective dog population management programs, HSI aims to reduce the size of free-roaming dog populations, improve their welfare, reduce shelter intake and euthanasia, as well as to reduce the number of accidental pet owners through the rehoming of unwanted puppies. Between 2013 and 2018, HSI in partnership with the Jamshedpur Utility Services Company Limited (JUSCO), a TATA Steel company appointed authority, implemented a CNVR program in Jamshedpur, India. During this program, free-roaming dogs were mostly hand caught on the street by trained animal handlers, vaccinated against rabies, neutered, and returned to the capture location. Between 2013 and 2016, the program neutered and vaccinated a total of 20,915 dogs. Street surveys were also conducted by HSI between 2014 and 2018 to measure the impact of this intervention on the free-roaming dog population size, structure, health, and welfare.

This study investigates whether HSI’s CNVR program in Jamshedpur was associated with changes in the health and welfare of the local free-roaming dog population, and the overall size, structure, and reproductive potential of the population. The objectives of this study are to: (i) determine whether, when comparing the same site over time, CNVR programs are associated with a reduction in the probability of (a) dogs entering the CNVR clinic with infectious and non-infectious clinical conditions, (b) dogs entering the CNVR clinic pregnant, (c) observing juvenile dogs in the free-roaming dog population (i.e. changing the structure of the population to older individuals), and, (d) observing lactating females in the free-roaming dog population; and (ii) determine whether the count of dogs observed in street surveys significantly declined throughout the study period.

## 2. Methods

### 2.1. Ethics Statement

Ethical review was not required for this study, as we analyse secondary data obtained from street dog sterilisation programs conducted according to standard operating procedures. No animals were specifically handled for the purpose of this study; instead, we analyse this dataset retrospectively to investigate the potential impact of government-supported animal birth control initiatives. In accordance with the Animal Welfare Board of India and the Animal Birth Control Rules of 2001, the Humane Society International has established written agreements with the local municipalities to carry out their dog sterilisation programs.

### 2.2. Study sites

This study investigates data collected by HSI as part of a CNVR programme conducted in Jamshedpur, India (Fig 1). Jamshedpur is the third biggest city in the state of Jharkhand, in north-eastern India. It has a tropical climate with mean daily temperatures between 12°C and 39°C and a wet season between June and October. The Greater Jamshedpur Metropolitan Region has a human population size of 1,339,438 (40). The CNVR project focussed on the JUSCO run township area (Jamshedpur Central). The study area was split into the following sites: Adityapur, Baridih, Bistupur, Golmuri, Jugsalai, Mango I, Mango II, Sidhgora, Sonari, and Telco. All sites had direct CNVR intervention, apart from Adityapur, Mango I, and Mango II, which acted retrospectively as negative controls. This CNVR project was launched in 2013, in partnership with the Animal Health Foundation (an animal welfare organisation) and ran until 2017.

**Fig 1.**
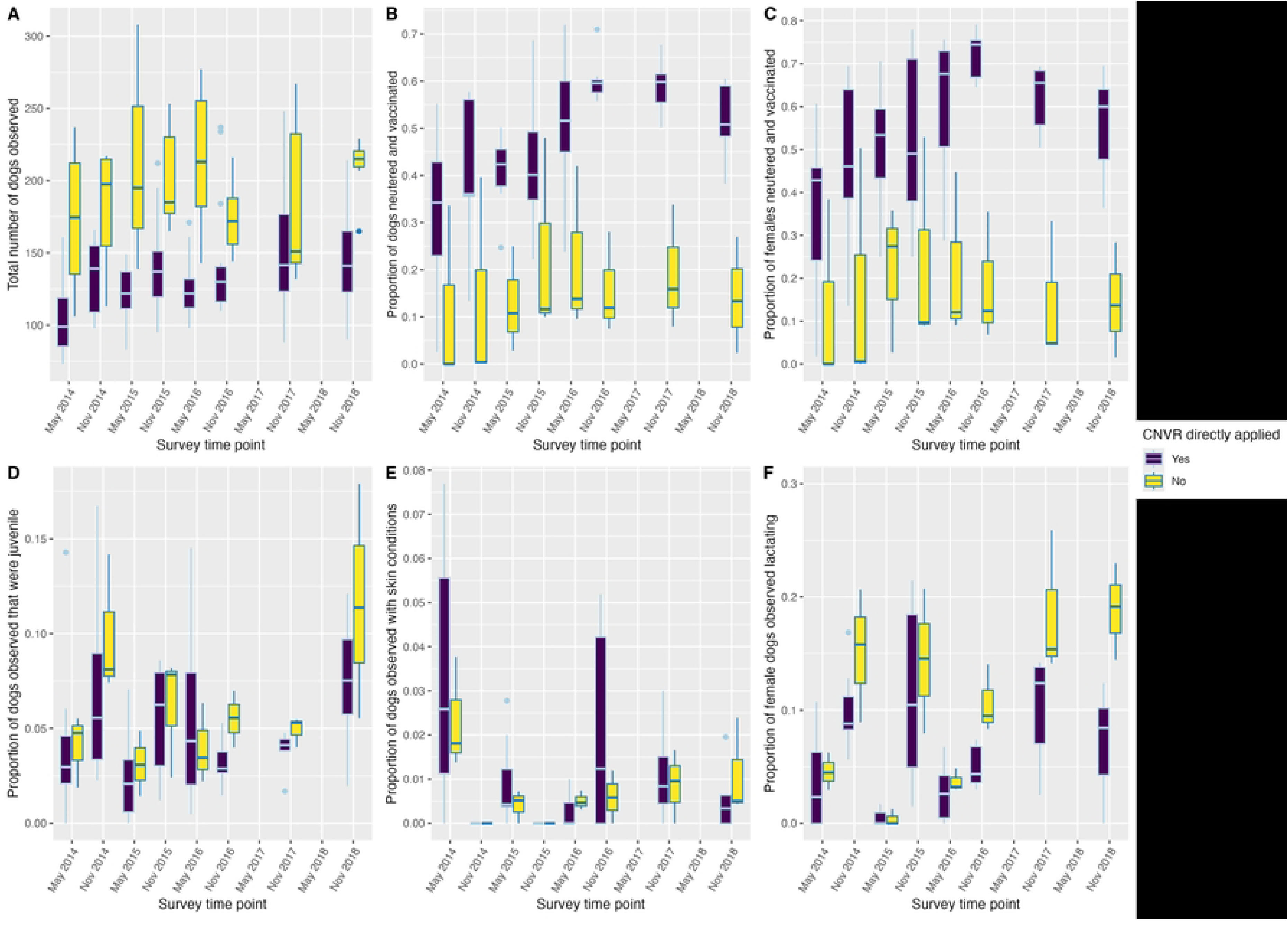
Inset map of India highlighting the state of Jharkhand and Jamshedpur. Map created using the package ggmap in R (42).

### 2.3. Data collection

Data on the free-roaming dog population was collected through street surveys conducted between May 2014 and December 2018. Clinical data was recorded from all dogs captured as part of the CNVR program between July 2013 and December 2016. Both clinical and street survey data were collected as part of the CNVR campaign by HSI staff members, and later shared with the authors for analyses.

#### 2.3.1. Street surveys

Street surveys were carried out bi-annually in May and November between 2014 and 2016, and then annually in November/December in 2017 and 2018. All study sites were surveyed at least once on each sampling occasion (i.e. no study sites had missing surveys). Within each study site, survey routes of approximately 25-30km in length were selected along existing pathways. Distances between study sites varied between 1 and 10 km. Surveys were conducted by two surveyors simultaneously using motorbikes and were usually conducted in the early morning. Surveyors travelled along the pre-determined routes moving at no more than 15km/hr. Every free-roaming dog was recorded, excluding those enclosed in private property or walking on a lead. At each sampling occasion, surveys were typically conducted twice over two consecutive days. However, there were instances where only one survey was conducted, accounting for 11% of the total site/sampling occasions, or three consecutive surveys were conducted, comprising 8% of the total site/sampling occasions.

Once a free-roaming dog was identified, information was collected by the surveyors using the mobile app OSM tracker to record each individual dog. Information on the dog’s age (adult; dogs that looked more than four months of age, or juvenile; dogs that looked four months or younger), sex (male/female), lactation status (lactating/not lactating), and neutering status (determined by presence of ear notch) were determined visually and recorded. In addition, the presence/absence of visible skin conditions was recorded, based on the presence/absence of hair loss, dermatitis, or visible ectoparasites. Body condition was recorded, based on a scale from one to five, relating to the dog’s body fat coverage (1 = emaciated, 2 = thin, 3 = ideal, 4 = overweight, 5 = obese) (41).

#### 2.3.2. Clinical data

Clinical data was collected by HSI veterinary clinic staff for all dogs captured as part of the CNVR program. All dogs were neutered and treated at a fixed clinic location. After dogs underwent surgery, they were kept in the clinic for a minimum of three days for post-operative care and monitoring. Clinical data included a description of the dog, how the dog was delivered to the clinic (from owner, caught by hand/net), date of operation, total length of stay in clinic, weight, reproductive status (in season/not in season; pregnant/not pregnant), treatment provided (vaccination; neutering; and Ivermectin administration), length of operation, and clinical conditions. We investigated whether time since implementation of CNVR was associated with changes in the probability of dogs entering the clinic with canine distemper virus, mange, rabies, canine transmissible venereal tumours and, for the female dogs, pregnancies. Rabies and canine distemper were diagnosed symptomatically, and transmissible venereal tumour disease and mange were diagnosed by visible appearance by qualified veterinarians at the clinic.

### 2.4. Statistical analyses

All statistical analyses were run in a Bayesian analysis framework using the package **brms** version 2.14.4 (43), in R version 4.3.1 (44). A negative binomial mixed effects model was fit to determine whether the number of dogs counted during street surveys decreased following CNVR intervention. Binomial mixed models were fit to assess whether the probabilities of observing: (i) sterilised dogs, (ii) juvenile dogs, or (iii) lactating females (for female subset of data) during street surveys decreased following CNVR intervention. To allow for overdispersion, we fit a beta-binomial model to assess whether the probability of observing dogs with skin conditions decreased following CNVR intervention. Models had demographic or health parameters as the outcome variables, and (i) survey number, (ii) CNVR (direct = study sites where dogs were caught, neutered, vaccinated, and released; non-direct = study sites where no/few dogs were caught, neutered, vaccinated, and released); and (iii) month (beginning of year (May) / end of year (November/December)) as predictor variables. Survey number is included so that May 2014 was survey one, November 2014 was survey two, May 2015 was survey three, and so on. Since only annual surveys were conducted in November/December in 2017 and 2018, the surveys normally scheduled for May (numbers seven and nine) are missing to account for the extended period between surveys. Survey site was included as a random intercept. As there may be different associations between the outcome variables for sites where CNVR was directly applied over time, we compared models with additive effects of predictor variables, and a model with an interactive effect between CNVR and survey time point. We determined the best fitting model using kfold cross-validation (10-folds) using the **kfold** function in the **brms** package (43). Kfold cross-validation involves subsetting the data into approximately equal folds. The model is then trained and assessed k-times using different folds as the test set to determine model fit.

To investigate whether the CNVR intervention was associated with changes in the probabilities of dogs entering the CNVR clinic with infectious conditions (canine distemper virus, mange, rabies virus, and canine transmissible venereal tumour disease), Bernoulli logistic regression models were fit with each of the infectious conditions as the outcome variables and year, month (to account for non-linear seasonal dynamics, we fitted using a generalised additive model function in the brms package), age (adult/juvenile, determined visually), and sex (male/female, determined visually) as predictor variables. To determine whether the CNVR intervention was associated with changes in the probabilities of females entering the CNVR clinic in a pregnant state, a Bernoulli logistic regression model was fitted on a subset of the data for females only, with pregnancy as the outcome variable and year, month (to account for non-linear seasonal dynamics, we fitted using a generalised additive model function in the **brms** package), and age (adult/juvenile) as predictor variables. As there may be different associations between the outcome variable for adults/juveniles across different sexes, for each of the clinical conditions, we compared models with both additive and interactive effects of age and sex (excluding pregnancy, as this analysis included a subset of the data for female dogs only). We also evaluated whether models were better fit including a seasonal trend by comparing models with and without month as a predictor variable. We determined the best fitting model using kfold cross-validation (10-folds).

All models were run with four chains, each with 2000 iterations (1000 used for warmup and 1000 for sampling). Thinning was set to one. The total number of post-warmup samples was 4000. Missing data was omitted from the statistical analysis (see supplementary information for details). All models had flat prior distributions. Model parameters were summarised by the mean and 95% credible intervals (CI; 95% most probable values). A significant effect was determined if the 95% credible intervals of the posterior distribution did not contain zero on the log odds or log scale. All predictor variables were centred, allowing for clearer interpretation of the results. Posterior distribution estimates were converted from logit scale to odds using exp (*x*), where *x* is the posterior value on the logit scale. All reported models converged (for all parameters Rhat ∼ 1.00 and effective sample size >1000, see Supplementary information).

## 3. Results

### 3.1. Street surveys – descriptive statistical analysis

Table 1 outlines the demographic factors and health indicators of dogs observed in street surveys between May 2014 and December 2018. The mean number of dogs counted during surveys (+/− SD) varied between 105 (+/−30) and 214 (+/− 52) (Table 1). In study sites where CNVR was directly applied (i.e. the seven sites where dogs were caught, neutered, and released), the percentage of sterilised dogs observed increased from 27% to a maximum of 61% across the study period. This provides our best estimate of CNVR coverage in this study. In the three study sites with no direct CNVR the percentage of sterilised dogs observed varied between 13% and 26% across the study period, with no consistent trend. The percentage of both lactating females and juveniles observed during surveys fluctuated annually with an increase in November and a decrease in May. The maximum percentage of juveniles observed was 8% in study sites where CNVR was directly applied, and 11% in sites where CNVR was not directly applied. The maximum percentage of females that were observed to be lactating was 11% in study sites where CNVR was directly applied, and 19% in sites where CNVR was not directly applied. Most dogs had a normal body condition score. Few dogs were observed with skin conditions throughout the study period. Fig 2 shows the number of dogs counted, the proportion of dogs observed that were neutered, that were female and neutered, female and lactating, adults, and observed with a skin condition across all time points for sites where CNVR was directly (i.e. sites where dogs were caught, neutered, vaccinated and release), and not directly applied (i.e. sites where dogs were not caught, neutered, vaccinated, and released).

**Fig 2.**
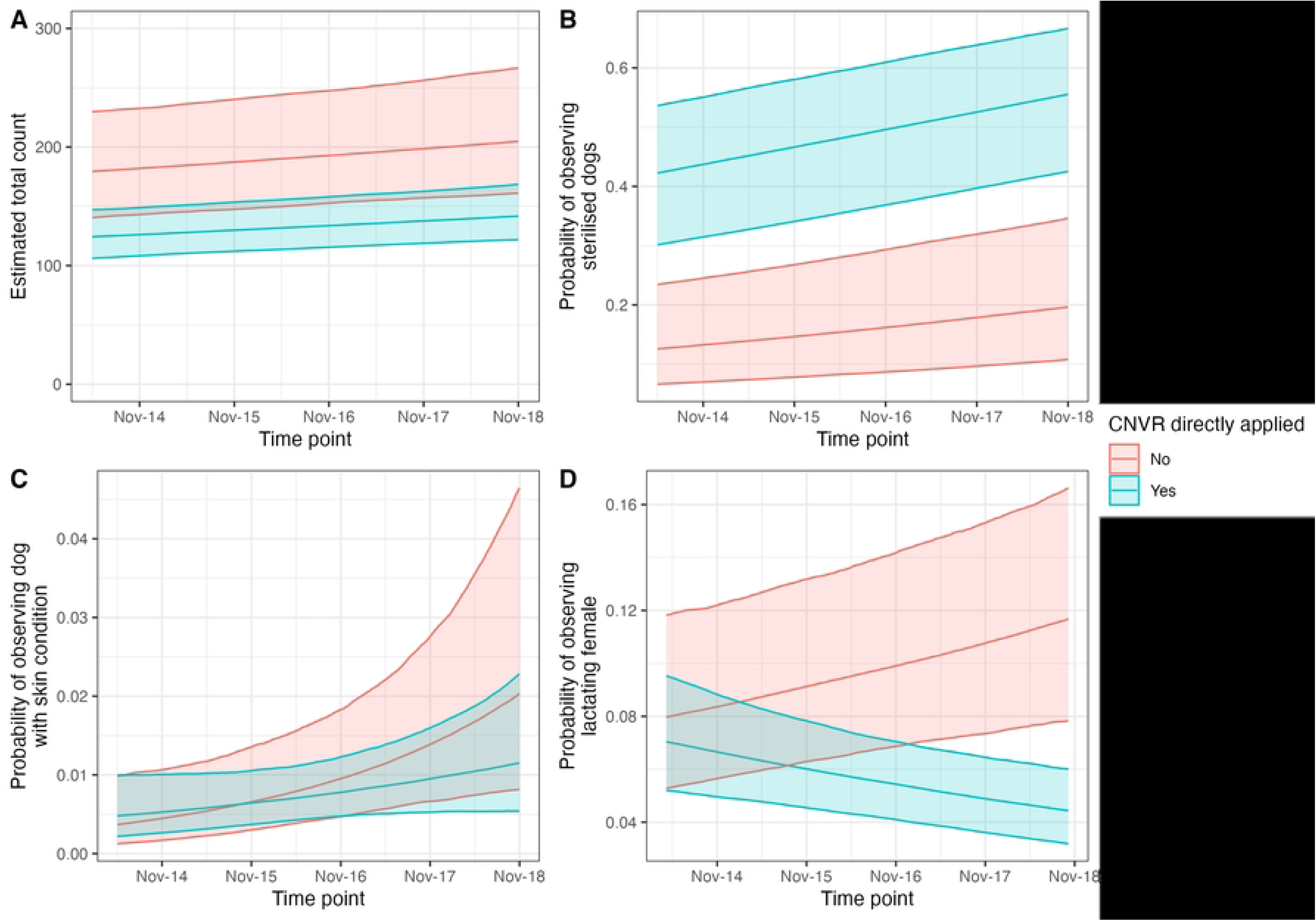
The total count of dogs observed during street surveys (A), and the proportion of dogs observed during street surveys that were neutered (B), female neutered (C), female lactating (D), juveniles (E), and observed with skin conditions (F) across survey period. Note: (i) that surveys were only conducted once per year in 2017 and 2018 in November/December and (ii) varying y-axis scales.

**Table 1.** Mean number (standard deviation; SD) of dogs counted during street surveys, and mean demographic and health factors of dogs observed between May 2014 and December 2018 for sites with direct CNVR (C; highlighted grey) and no direct CNVR (N) applied.

|  | C | N | C | N | C | N | C | N | C | N | C | N | C | N | C | N | C | N | C | N | C | N |
| --- | --- | --- | --- | --- | --- | --- | --- | --- | --- | --- | --- | --- | --- | --- | --- | --- | --- | --- | --- | --- | --- | --- |
| Time point | Mean count (SD) | Mean count (SD) | Sterilised (%) |  | Females sterilised (%) |  | Juvenile (%) |  | Skin Conditions (%) |  | Females lactating (%) |  | Body condition score |  |  |  |  |  |  |  |  |  |
|  |  |  |  |  |  |  |  |  |  |  |  |  | 1 |  | 2 |  | 3 |  | 4 |  | 5 |  |
| May-14 | 105 (30) | 173 (59) | 27 | 21 | 29 | 25 | 4 | 4 | 4 | 2 | 5 | 4 | 1 | 0 | 0 | 0 | 95 | 98 | 2 | 1 | 1 | 0 |
| Nov-14 | 132 (26) | 181 (43) | 42 | 16 | 48 | 19 | 7 | 10 | 0 | 0 | 10 | 15 | 0 | 0 | 0 | 0 | 100 | 100 | 0 | 0 | 0 | 0 |
| May-15 | 121 (20) | 214 (86) | 41 | 16 | 52 | 26 | 2 | 4 | 1 | 0 | 1 | 1 | 0 | 0 | 45 | 38 | 55 | 61 | 0 | 0 | 0 | 0 |
| Nov-15 | 142 (32) | 201 (37) | 40 | 26 | 49 | 30 | 6 | 6 | 0 | 0 | 11 | 13 | 0 | 0 | 0 | 0 | 100 | 100 | 0 | 0 | 0 | 0 |
| May-16 | 125 (21) | 214 (52) | 50 | 24 | 60 | 24 | 6 | 4 | 0 | 1 | 3 | 4 | 0 | 0 | 0 | 0 | 99 | 99 | 0 | 0 | 0 | 0 |
| Nov-16 | 144 (43) | 175 (27) | 61 | 20 | 73 | 24 | 3 | 5 | 3 | 1 | 6 | 10 | 0 | 0 | 1 | 0 | 99 | 100 | 1 | 0 | 0 | 0 |
| Nov-17 | 150 (42) | 184 (61) | 58 | 22 | 63 | 18 | 4 | 5 | 1 | 1 | 10 | 18 | 0 | 0 | 0 | 0 | 100 | 100 | 0 | 0 | 0 | 0 |
| Dec-18 | 144 (35) | 210 (21) | 50 | 13 | 53 | 14 | 8 | 11 | 0 | 1 | 6 | 19 | 0 | 0 | 0 | 0 | 100 | 100 | 0 | 0 | 0 | 0 |

### 3.2. Street surveys – inferential statistical analysis

#### 3.2.1. Total count of dogs observed

The model with an additive effect between CNVR and survey timepoint was a better fit than an additive model (Table S1). Sites where CNVR had been directly applied had significantly lower counts of dogs (rate ratio 0.69, 95% CI 0.51-0.90). In sites where CNVR had been directly applied, the average model estimated count of dogs was 133 (95% CI 113-153), compared to 193 (95% CI 149-242) in areas where no direct CNVR was applied. The total count of dogs significantly increased over time (rate ratio 1.01, 95% CI 1.00-1.03; Fig 3A). There was no evidence of a significant effect of biannual survey month (May versus November/December) (rate ratio 1.01, 95% CI 0.998-1.021).

**Fig 3.**
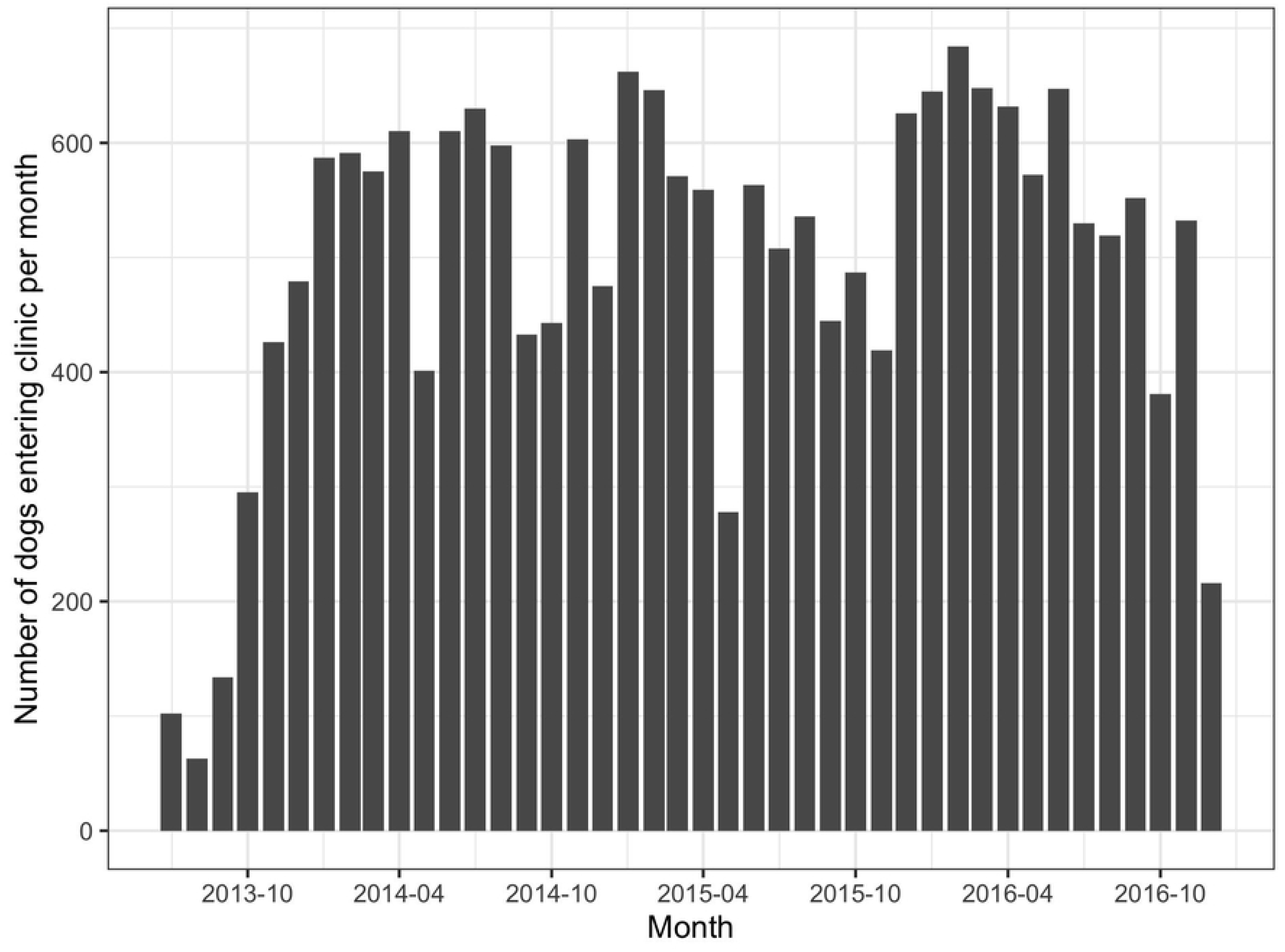
(A) Model estimated total count of dogs observed over time, (B) estimated probability of observing sterilised dogs over time, (C) observing a dog with a skin condition, and (D) observing lactating female for sites with CNVR directly applied and not. Ribbons show the 2.5 and 97.5 percentiles of the posterior distribution (95% CI).

#### 3.2.2. Proportion of sterilised dogs

The model with an additive effect between CNVR and survey timepoint was a better fit than an interactive effect (Table S1). The model estimated probability that an observed dog was sterilised significantly increased over time (OR 1.06, 95% CI 1.05-1.07; Fig 3), and there was a significantly higher probability of observing sterilised dogs in sites where CNVR had been directly applied (OR 5.64, 95% CI 1.59-11.00). The estimated probability of observing a sterilised dog in sites with direct CNVR was 0.46 (95% CI 0.35-0.57), compared to 0.15 (95% CI 0.05-0.16) in sites without direct CNVR. There was no evidence of significant associations between biannual survey month (May versus November/December) and the probability of observing sterilised dogs (OR 1.01, 95% CI 0.99-1.02).

#### 3.2.3. Proportion of juvenile dogs

The model with an additive effect between CNVR and survey timepoint was a better fit than an interactive model (Table S1). The month of the survey (May or November/December) was significantly associated with the probability that an observed dog was juvenile (OR 1.07, 95% CI 1.04-1.10). In May, the model estimated probability that an observed dog was juvenile (0.04, 95% CI 0.03-0.5) was significantly lower than in November/December (0.06, 95% CI 0.05-0.07). There was no evidence of a significant association between the estimated probability that an observed dog was juvenile and survey time point (OR 1.02, 95% CI 0.999-1.042) or sites where CNVR had been applied directly (OR 0.77, 95% CI 0.51-1.04).

#### 3.2.4. Proportion of lactating females

The model with an interactive effect between CNVR and survey timepoint was a better fit than an additive model (Table S1). The month of the survey (May or November) was significantly associated with the probability that an observed female dog was lactating (OR 1.29, 95% CI 1.23-1.35). In May, the model estimated probability that an observed female dog was lactating (0.03, 95% CI 0.02-0.03) was significantly lower than in November/December (0.11, 95% CI: 0.09-0.13). There was a significant interactive effect of CNVR and time point (OR 0.90, 95% CI 0.86-0.95; Fig 3D).

#### 3.2.5. Proportion of dogs with skin conditions

The model with an additive effect between CNVR and survey timepoint was a better fit than an interactive effect (Table S1). The biannual survey month (May versus November/December) was significantly associated with the probability that an observed dog had a skin condition (OR 0.85, 95% CI 0.77-0.93). In May, the model estimated probability that an observed dog had a skin condition (0.02, 95% CI 0.009-0.025) was significantly higher than in November/December (0.007, 95% CI 0.004-0.011). The probability that an observed dog had a skin condition significantly increased over time (Fig 3C; OR 1.14, 95% CI 1.03-1.27). There was no evidence of significant associations between observing dogs with skin conditions and sites where CNVR had been directly applied (OR 0.87, 95% CI 0.27-3.51).

### 3.3. Clinical data – descriptive statistical analysis

In total, 20,915 dogs were taken into the clinic to be neutered and vaccinated in the CNVR program in Jamshedpur between 4^th^ July 2013 and 13^th^ December 2016. A mean of 498 (sd +/− 155) dogs were neutered and vaccinated every month (Figure 4). Of these 13,062 (62%) were caught by hand, 7,448 (36%) were caught by net, 305 (1%) were brought in by owners, 85 (0.4%) were brought into the clinic by unreported method (missing information), and 15 (0.07%) were trapped. In total, the CNVR program neutered 10,459 (50%) females, 10,218 (49%) male dogs, and 238 (1%) dogs with missing information on their sex. The CNVR program neutered 13,087 (63%) adult dogs, 7,590 (36%) juvenile dogs, and 238 (1%) dogs with missing information on their age. Figure 5 shows the percentages of dogs per month that were diagnosed by visible appearance as positive for canine distemper virus, mange, rabies, and canine transmissible venereal tumour. Figure 6 shows the percentages of female dogs per month that entered the clinic pregnant; as previously reported the percentages of pregnant dogs peaked each year in September/October (45).

**Figure 4.**
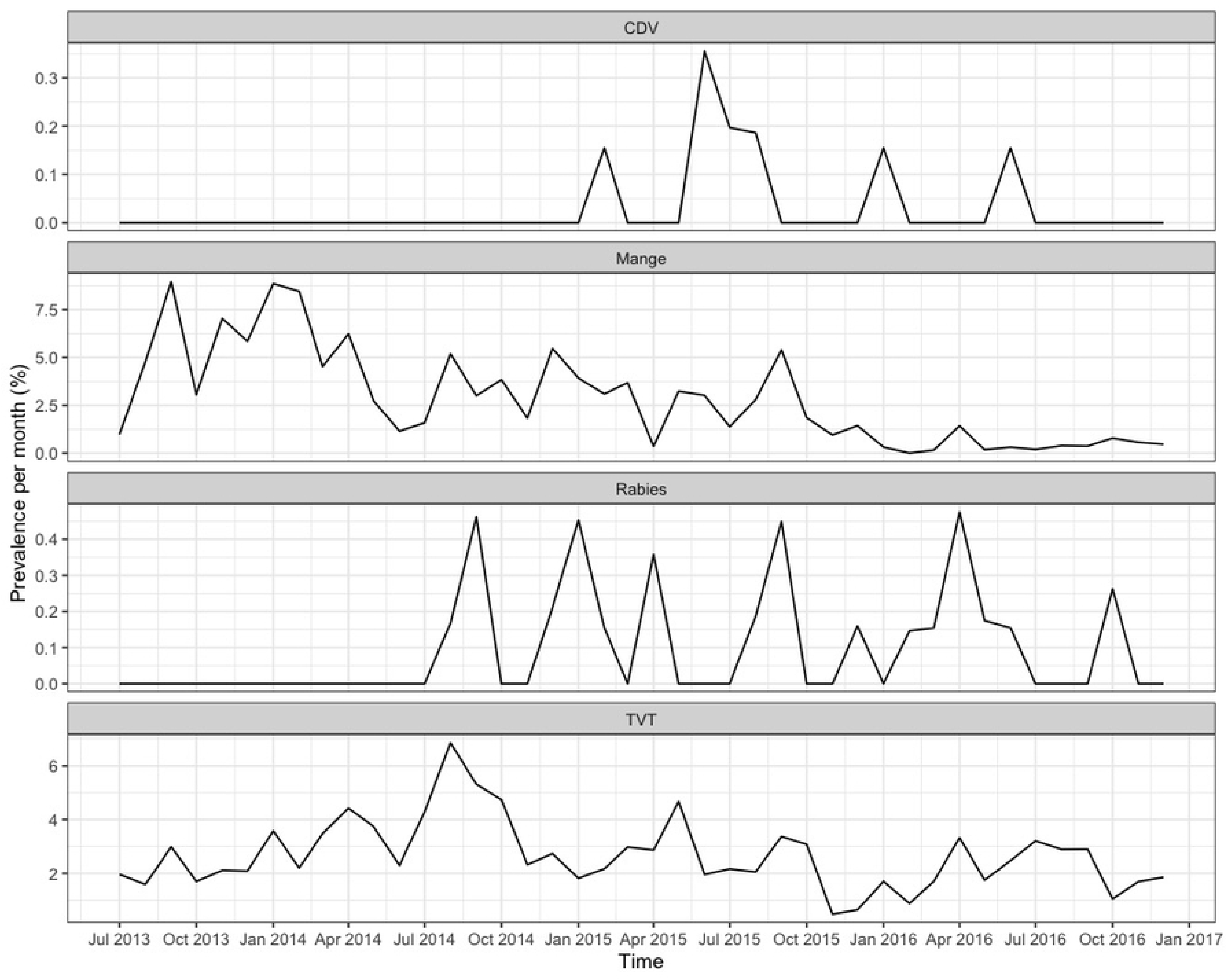
Number of dogs entering clinic per month as part of the Humane Society International’s Catch-Neuter-Vaccinate-Release intervention in Jamshedpur, India.

**Figure 5.**
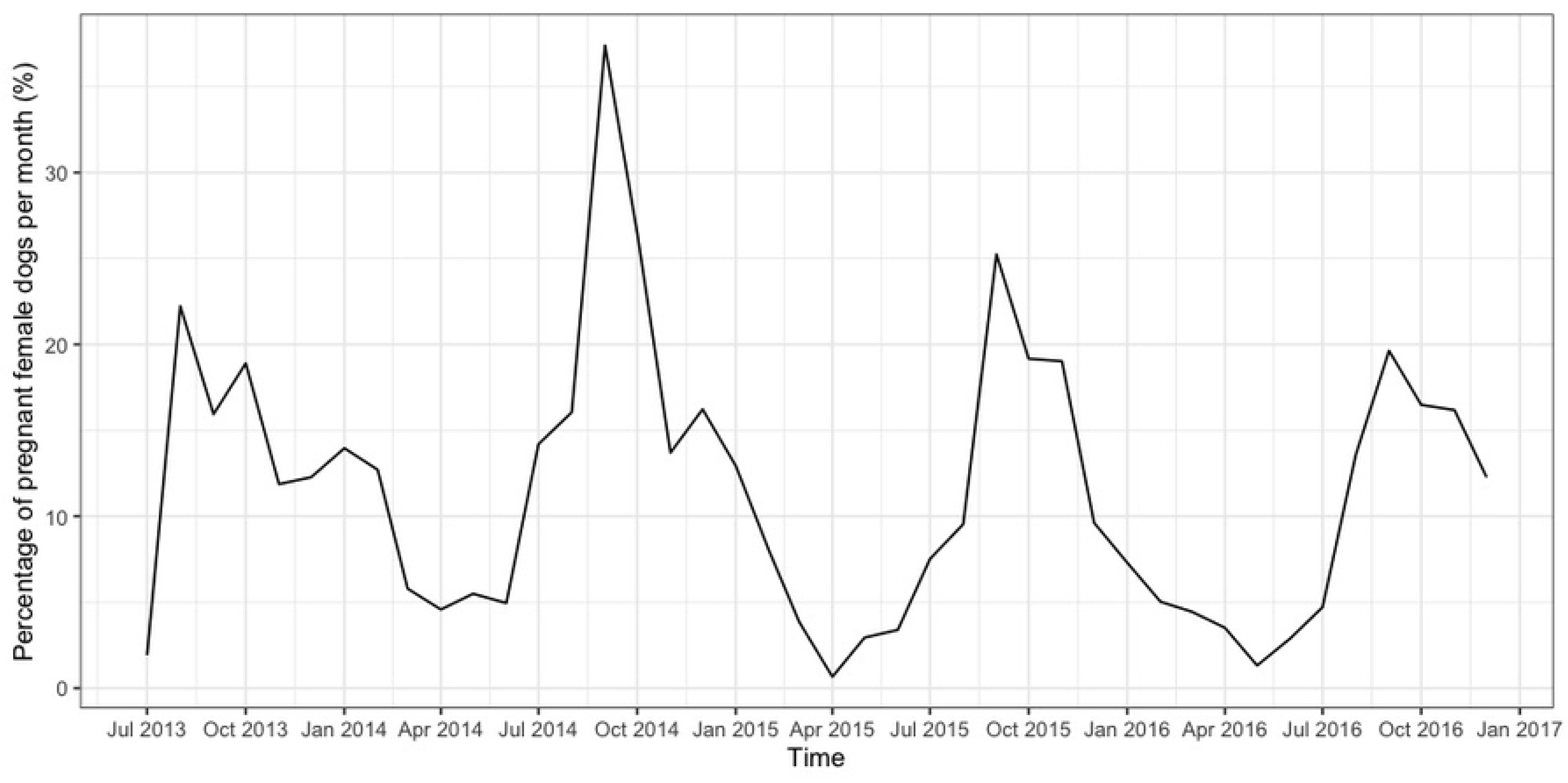
The percentages of dogs per month in Jamshedpur CNVR clinic that were diagnosed by visible appearance as positive for canine distemper virus (CDV), mange, rabies virus, and canine transmissible venereal tumour (TVT). Note the different y-axis scales per plot.

**Figure 6.**
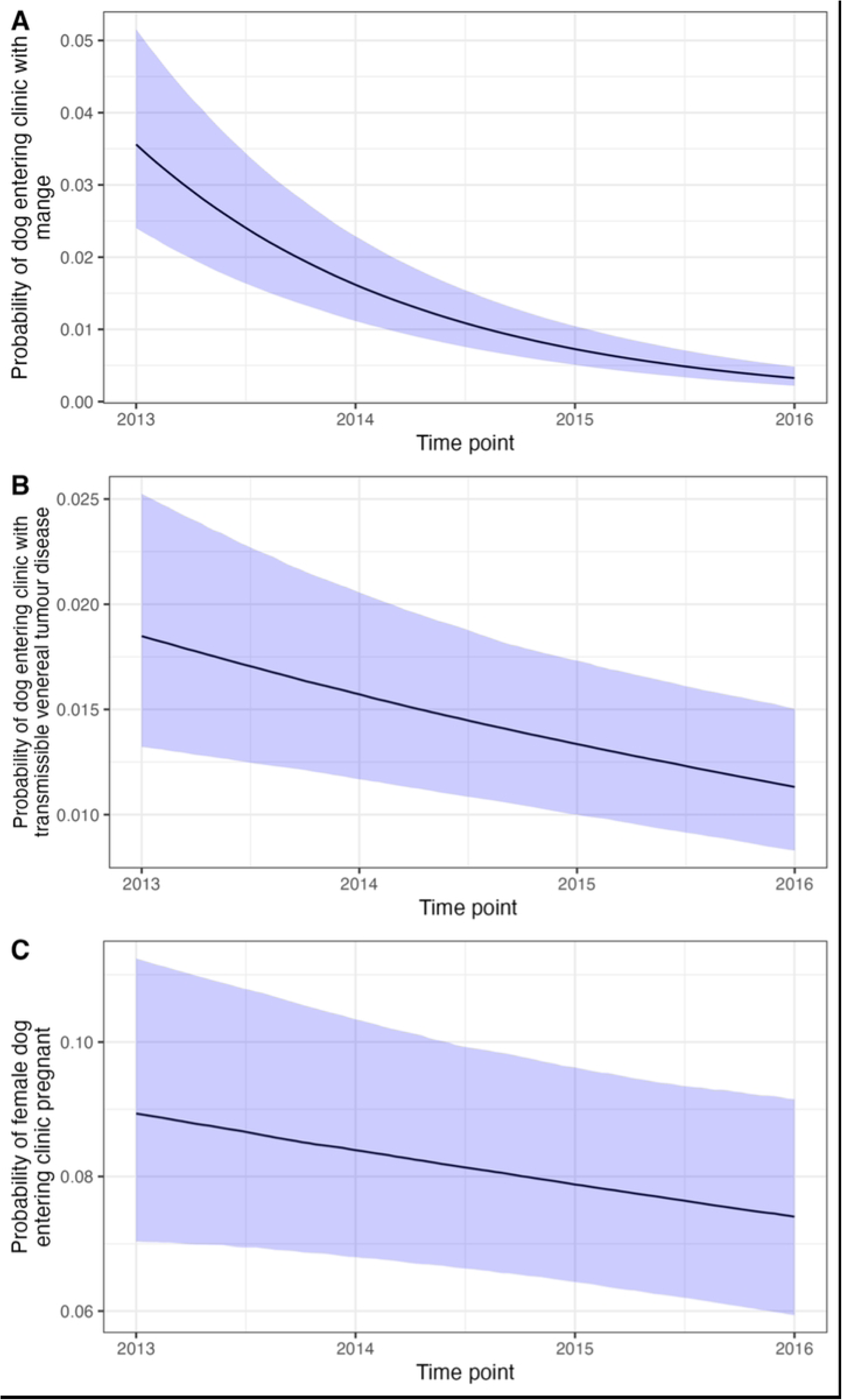
The percentages of female dogs per month entering the clinic in Jamshedpur CNVR clinic that were pregnant.

### 3.4. Clinical data – inferential statistical analysis

#### 3.4.1. Proportion of dogs entering clinic with mange

The model with month of the year included and age and sex as additive fixed effects was best fitting (Table S1). The model estimated probability of dogs entering the clinic with mange significantly decreased with year since the start of the intervention (OR 0.45, 95% CI 0.40-0.50; Figure 7). There was a significant effect of month (mean and 95% credible intervals = 0.13, 95% CI 0.09-0.16; Figure 8). Female dogs (OR 0.63, 95% CI 0.52-0.74) had a significantly lower probability of entering the clinic with mange, female dogs had a 1.29% (95% CI 1-1.41% probability compared to 1.90% (95% CI 1.66-2.20%) for males. Adult dogs (OR 4.32, 95% CI 3.22-5.55) had a significantly higher probability of entering the clinic with mange, adult dogs had a 2.54% (95% CI 2.22-2.82%) probability compared to 0.61 (95% CI 0.46-0.78%) for juveniles.

**Figure 7.**
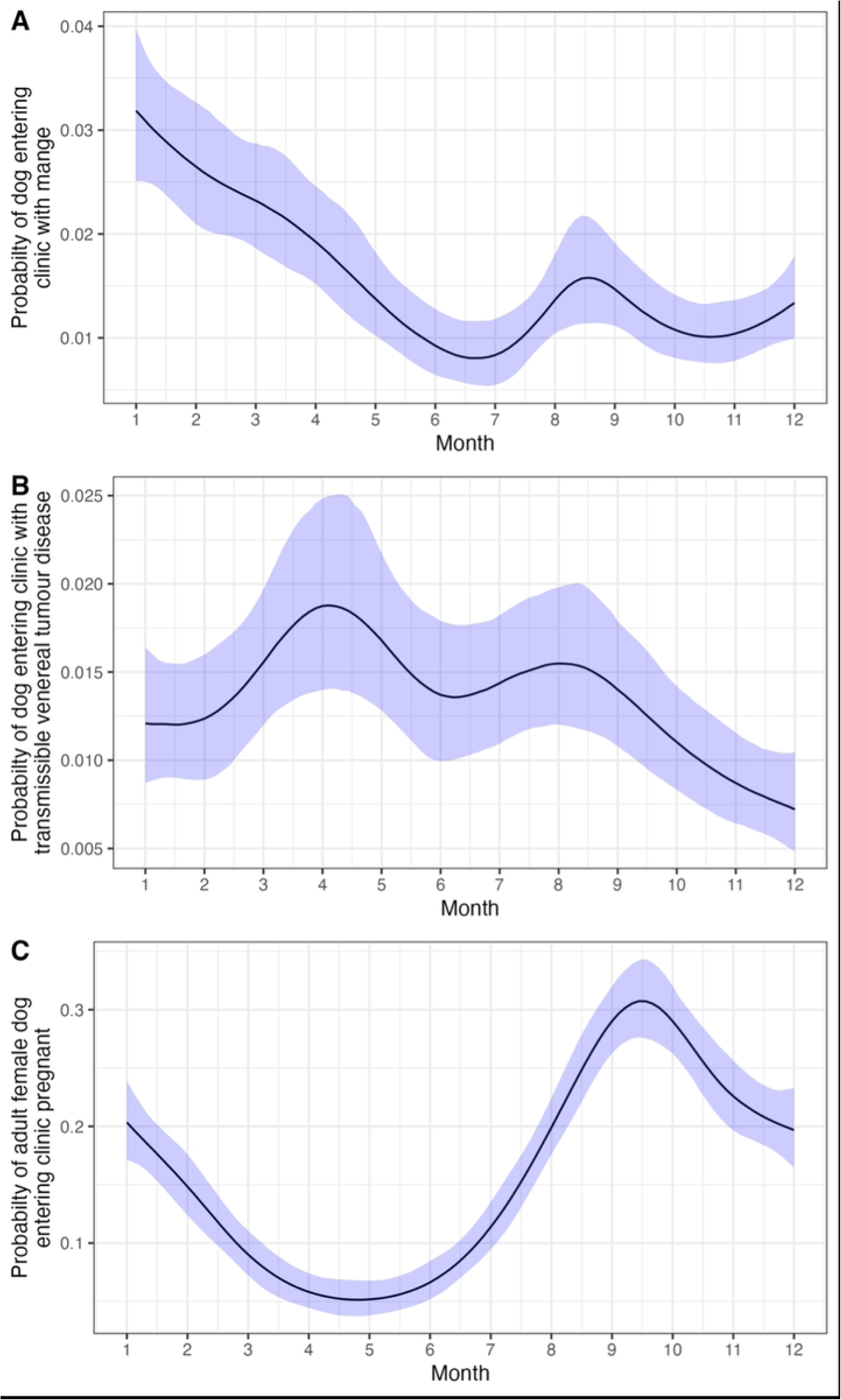
Statistical association between year and the probability of a dog entering the clinic with (A) mange, (B) transmissible venereal tumour disease, and (C) pregnant (female only). Estimated probabilities are averaged for all other predictor variables, apart from pregnancy, which reports the estimated effect for an adult dog. Ribbons show the 2.5 and 97.5 percentiles of the posterior distribution (95% CI).

**Figure 8.**
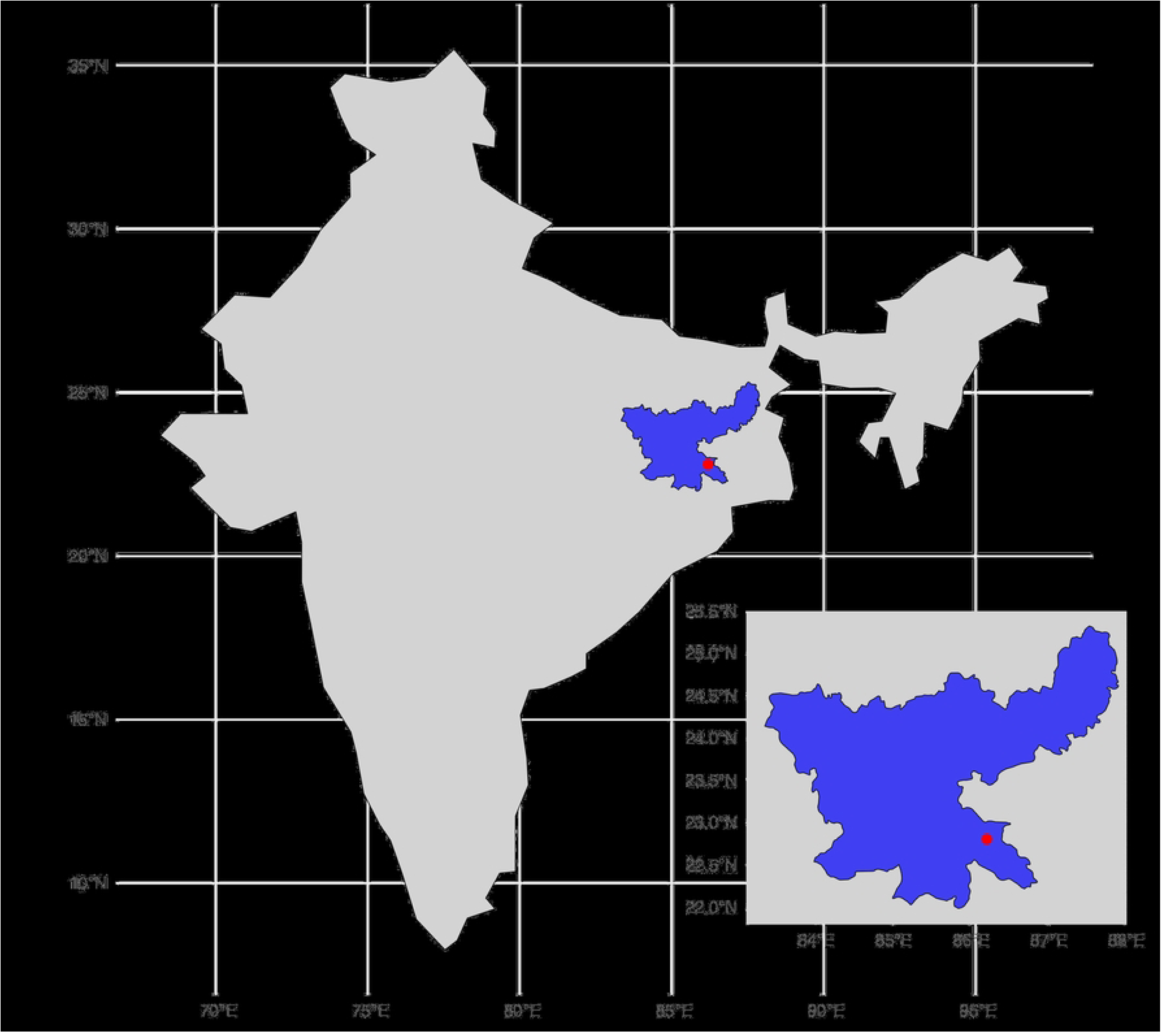
Statistical association between month and the probability of a dog entering the clinic with (A) mange, (B) transmissible venereal tumour disease, and (C) pregnant (female only). Estimated probabilities are centred for all other predictor variables, apart from the association between month and pregnancy, which reports the estimated effect for an adult dog. Ribbons show the 2.5 and 97.5 percentiles of the posterior distribution (95% CI).

#### 3.4.2. Proportion of dogs entering clinic with canine transmissible venereal tumour

The additive model with a seasonal component was best fitting (Table S1). The model estimated probability of dogs entering the clinic with canine transmissible venereal tumours significantly decreased with the year (OR 0.85, 95% CI 0.77-0.94; Figure 7). Adult dogs had a higher estimated probability of entering the clinic with canine transmissible venereal tumour (OR 16.24, 95% CI 9.30-24.05), adult dogs had a 3.44% (95% CI 3.11%-3.77%) probability of entering the clinic with canine transmissible venereal tumours, compared to 0.23% (95% CI 0.13%-0.33%) for juvenile dogs. Female dogs (OR 2.49, 95% CI 2.05-2.98) had a higher probability of entering the clinic with canine transmissible venereal tumours, female dogs had a 1.99% (95% CI 1.65%-2.36%) probability of entering the clinic with canine transmissible venereal tumours, compared to 0.82% (95% CI 0.63%-0.99%) for male dogs. The probability of dogs entering the clinic with canine transmissible venereal tumour disease was statistically significantly associated with the month of the year (mean and 95% credible intervals = 0.07, 95% CI 0.03-0.11; Figure 8).

#### 3.4.3. Proportion of dogs entering clinic with rabies

The model with the interactive effect of sex and age, without a seasonal component was best fitting (Table S1). There were no significant statistical associations between the probability of dogs entering the clinic with rabies and year (OR 1.93, 95% CI 0.82-3.18), age (OR 1.03, 95% CI 0.16-2.39), sex (OR 1.12, 95% CI 0.18-2.55), or interactive effect of age and sex (OR 13.20, 95% CI 0.09-44.22).

#### 3.4.4. Proportion of dogs entering clinic with canine distemper virus

The model with an additive effect of sex and age, without a seasonal component was best fitting (Table S1). There were no significant statistical associations between the probability of dogs entering the clinic with canine distemper virus and year (OR 1.89, 95% CI 0.43-4.19), age (OR 1.91, 95% CI 0.04-6.26), or sex (OR 20.58, 95% CI 0.13-57.79).

#### 3.4.5. Proportion of female dogs entering clinic pregnant

The model with month of the year included was best fitting (Table S1). The probability of female dogs entering the clinic pregnant significantly decreased with time since the start of the intervention (Figure 7; OR 0.93, 95% CI 0.87-0.99). Adult dogs had a higher probability of entering the clinic pregnant (OR 33.25, 95% CI 20.49-47.23), adult females had a 13.42% (95% CI 12.56-14.35%) probability of entering the clinic pregnant, compared to 0.48% (95% CI 0.30-0.69%) for juveniles. There was a significant association between the month of the year and probability of entering the clinic pregnant (mean and 95% credible intervals = 0.137, 95% CI 0.109-0.167; Figure 8).

## 4. Discussion

This study determined whether the introduction of the CNVR program was associated with changes in (i) the health and welfare of the local free-roaming dog population, and (ii) the population’s overall size, structure, and reproductive potential. We found that the probability of dogs entering the clinic with mange, canine transmissible venereal tumour disease, or pregnant significantly decreased with time since the start of the CNVR programme. In the street surveys of free-roaming dogs, the probability of observing sterilised dogs increased over time, and there was a higher probability of observing sterilised dogs in sites where CNVR had been directly applied. Despite the overall increase in sterilised dogs, survey counts significantly increased over time, although lower counts of dogs were observed in CNVR-treated sites (Fig 3). The age-structure observed in street surveys did not significantly change over time. In CNVR-treated sites, the probability of observing lactating females declined, while in non-treated sites it increased (Fig 3).

### Health and Welfare

Most dogs observed in the street surveys had normal body condition scores and few dogs had observable skin conditions (Table 1, Fig 2). These results are similar to those reported by other studies in South Asia (46,47), although different to Totton *et al.* (30), who reported a high prevalence of dogs with low body conditions (70%) and skin conditions (69%) in Jodhpur, India. Free-roaming dogs are sustained from food primarily provided from people, either directly through deliberate feeding, or indirectly through human waste (22,48,49). The normal body condition scores reported in this study suggest sufficient supply of food for this free-roaming dog population. Bhalla *et al.* (22) reported that bakeries, houses, and garbage piles are significant sources of food to free-roaming dogs in Bangalore, India.

We observed a small, but significant positive correlation between the probability of observing dogs with skin conditions and time since the beginning of the CNVR intervention (Fig 3C), although no significant difference between CNVR versus non-CNVR treated sites. Previous research has reported an increase in the prevalence of skin conditions following CNVR interventions (29,30). This is possibly due to the conditions dogs are kept in the clinic both pre– and post-sterilisation, such as increased density of dogs that could facilitate the spread of infectious diseases, and whether antiparasitic treatment is administered. In this study, all dogs sterilised as part of the CNVR intervention were treated with Ivermectin, an antiparasitic drug. In the clinical analysis, we observed a significant decrease in the probability of dogs entering the CNVR clinic with mange following time since the beginning of the CNVR intervention (Figure 7), indicating a possible decrease in the prevalence of mange in the wider free-roaming dog population. The differences in estimated prevalence of skin conditions between these two datasets could relate to the different methods; dogs observed at distance in the street, versus close contact in the clinic. Mange is a skin disease caused by infection with parasitic mites and is one possible cause of observable skin conditions in free-roaming dog populations. The administration of Ivermectin to all dogs sterilised in the CNVR program could influence the dynamics of parasitic diseases in the wider free-roaming dog population (29,30), although further research would be required to ascertain this relationship.

We observed a significant decrease in the probability of dogs entering the clinic with canine transmissible venereal tumour disease following time since the beginning of the CNVR intervention (Figure 7). Canine transmissible venereal tumour disease is usually transmitted during mating and is one of the few naturally transmissible cancers in mammals. It has an estimated global prevalence of around 1% (50), but may be as high as 15% in females in some free-roaming dog populations (51). As sexually intact dogs mate more frequently, they are at higher risk of contracting this disease. If more dogs in the population are sterilised, the frequency of mating may reduce, therefore reducing the risk of exposure to this disease. We estimated that female dogs had a higher probability of canine transmissible venereal tumour infection (1.99%, 95% CI 1.65%, 2.36%), compared to males (0.82%, 95% CI 0.63%, 0.99%). This has been reported elsewhere and has been associated with the mating dynamics of free-roaming dogs (52,53); males are sexually receptive all year and one infected male often mates with several females, leading to a higher prevalence in female dogs (54,55).

We did not observe a significant association between time since the beginning of the CNVR program and the probability of dogs entering the clinic with rabies or canine distemper virus. In general, the monthly prevalence of dogs observed in the clinic with either distemper or rabies was low, therefore there was low power to determine statistically significant associations. It is also worth noting that dogs were diagnosed symptomatically, limiting our confidence in the reported estimates. As dogs were vaccinated against rabies as part of the CNVR program, we may have expected to observe a decline in the number of dogs entering the clinic with rabies. It is estimated that 70% vaccination coverage of dog populations is required to prevent a rabies disease outbreak (56,57); declines in canine and human rabies cases have been reported in areas following annual vaccination of 70% of the dog population (58,59). Given the neutering and vaccination coverage was consistently below 60% in the study sites (Fig 2B), a greater number of dogs may need to be vaccinated to observe a decline. Immunity towards rabies may also be lower, due to effects of waning vaccination if dogs are not regularly vaccinated.

The sampling of dogs taken into the clinic for surgical sterilisation represents a convenience sample, and here we assume that this sample is representative of the wider population. In reality, only intact dogs (or those assumed to be intact) and those able to be captured are included in this convenience sample. This may bias our estimated prevalence of infectious diseases in the wider free-roaming dog population. For example, dogs with rabies may be less likely to be caught, leading to underestimation of the prevalence of rabies in the wider dog population. A cross-sectional sero-prevalence survey of the dog population for antibodies against infectious diseases of interest may provide less-biased estimates of associations between CNVR programs and the prevalence of infectious diseases.

### Population size, structure, and reproductive potential

The counts of dogs observed in the street significantly increased over time (Fig 3A). This is possibly due to the low neutering coverages observed, which were often observed to be lower than 60% (Fig 2B). CNVR aims to decrease the birth rate in the free-roaming dog population, with the hypothesis that it will result in stabilisation, eventually leading to a reduction in dog population size. Neutering coverages maintained at 65% and higher over several years have been associated with declines in free-roaming dog populations in Jaipur, Jodhpur and Ranchi in India (26,36,37). Additionally, the human population size in Jamshedpur grew by approximately 2% annually, which may have increased resources for dogs or led to their migration, as dog population size often correlates to human population size.

If CNVR impacts dog population size, we might expect age structure changes prior to observing population decline. We observed no significant change in adult and juvenile proportions over time, including in CNVR-treated sites. This might be due to the low sterilisation coverage. We observed a small difference in age structure depending on the month of the survey: 6% (95% CI 5-7%) of the dogs observed in street surveys in November were juveniles, compared to 4% (95% CI 3-5%) in May. Although dogs generally do not exhibit seasonal breeding, in India, dogs have one clear breeding season (45,60). This season begins after the rainy season and high proportions of puppies are often observed at the end/beginning of the year. The age structures observed in this study align with those in Jodhpur and Ranchi (26,36), though we observed fewer juveniles that in Maharashtra, India (61,62), where juveniles comprised 15-30% of the population. These variations may reflect differences in CNVR interventions, or in visual age estimation methods. Juvenile dogs, particularly young puppies, may have a lower detection probability. For example, puppies may be hidden and less likely to be observed during street surveys, meaning proportions of adults and juveniles observed during street surveys might not reflect the true age structure in the population, possibly making it challenging to observe changes in the age structure over time.

Although we did not observe a decline in the numbers of dogs counted, or a change in the age structure, we observed a significant decrease in the probability of female dogs entering the clinic pregnant with time since the start of the intervention (Figure 7). If CNVR has reduced the birth rate, we may expect to see fewer pregnant females in the clinic if there are fewer intact dogs in the population. We also observed an interactive effect between time since the beginning of the intervention, whether CNVR was directly applied and the probability of observing lactating females (Fig 3). The probability of observing lactating females increased in areas where CNVR was not directly applied, compared to the probability decreasing in areas with direct CNVR intervention. Perhaps over a longer time-frame and with neutering coverage maintained above a critical threshold, we would observe a reduction in the free-roaming dog population size also.

### Study limitations

In this study, we compared population counts observed during street surveys through time. Simple count methods do not provide an estimate of the total population size, as this method does not account for imperfect detection of individuals. Instead, simple counts may act as an indicator of population size, showing relative trends over time. Simple count methods are logistically easier but their usefulness in tracking trends depends on whether they are directly proportional to the true population size (63,64). This means that the probability of detecting individuals must be constant across survey time points. It is unclear if this assumption is true for free-roaming dog populations. Variation in detection probability has been associated with the time of day, day of the week, and time of year (47,65,66). This may be controlled by ensuring surveys are conducted across consistent time, days, and months. Variation in detection probability due to other factors has been reported, such as days with food markets or festivals (66,67), and environmental conditions, such as wind velocity and ambient temperature (61). Estimates of population size using mark-resight techniques would avoid these issues, and allow organisations involved in dog population management to plan their interventions to ensure high neutering and vaccination coverage.

Additionally, although we compared sites with and without directly applied CNVR, neutered dogs were observed in both locations. This suggests movement of dogs between sites, and although lower numbers of neutered dogs were observed in sites without CNVR (Fig 2), this will have reduced our ability to detect an effect of CNVR. Study designs, such as stepped-wedge interventions, where interventions are applied systematically to areas over time, with each area acting as a control at the beginning may be beneficial for future investigating the impact of dog population management interventions. This study design also overcomes the ethical challenge of keeping some areas as controls.

## 5. Conclusions and future recommendations

Over recent years, there has been increasing appreciation of the importance of measuring the impact of existing dog population management programmes (27,68). This is important for gaining insight into the factors that influence the success of management strategies. The information presented in this study contributes to this growing body of research. Our results suggest that a low coverage of around 60% may be associated with a reduction in mange and canine transmissible venereal tumour disease, and a reduction in the proportion of lactating females, but is not enough to reduce the population size. This knowledge can help guide future dog population management efforts, suggesting organisations should aim to maintain high sterilisation coverages. Further monitoring and evaluation of CNVR program impacts are required to determine optimal coverages required to reduce population size.

We observed clear seasonality in our study sites in Jamshedpur. This suggests that to optimise CNVR interventions, maximising sterilisation coverage prior to the breeding season could potentially allow efficient and effective high intervention coverages to be maintained during the critical breeding period. This approach may reduce the resources and costs required, such as staffing and facilities. The impact of timing CNVR interventions with the seasonal breeding cycle of Indian free-roaming dogs requires further investigation. For example, modelling the effect of timing interventions prior to the breeding season on intervention costs and population size.

## Acknowledgements

We extend our gratitude to Dr Andrew Rowan for initiating this collaboration and for sharing his important knowledge and input into this project. We also wish to thank the dedicated staff at the Humane Society International, whose many hours of effort in project management, implementation, data collection, and collation were invaluable to this project.

